# Coordinated Drift of Image Representations in Human EEG

**DOI:** 10.64898/2026.10.27.762314

**Authors:** Yile Wang, Zitong Lu, Kaushik Lakshminarasimhan

## Abstract

Sensory representations drift over time in many animal systems, but evidence in humans remains limited. Moreover, how perception remains stable despite representational drift remains unclear. Here, we used THINGS-EEG2, a dataset with multiple sessions of repeated presentations of natural images, to test whether image-evoked EEG responses drift across recording sessions and whether this drift is geometrically constrained. We found that image-specific EEG response patterns drifted in a manner that was coordinated across channels and concentrated along dimensions that encoded image identity. Although this impaired cross-session decoding accuracy on average, its effects were heterogeneous with some image pairs becoming harder to distinguish and others becoming more separable. Critically, global distance statistics and the dimensionality of the image response space were preserved across sessions. Simulations show that downstream Hebbian networks can adapt faster to structured drift that preserves dimensionality than when drift is unstructured. These findings suggest that human sensory representations can change over time while remaining amenable to rapid compensation via unsupervised learning mechanisms.

## 2 Introduction

We can instantly recognize a familiar face or a favorite painting. This perceptual constancy might suggest that the sensory representations that drive those percepts are themselves stable over time. However, longitudinal neural recordings in many brain areas have shown that neural response patterns change substantially over time, even when stimuli and behavioral demands remain unchanged (Chambers et al., 2023; Driscoll et al., 2022; Rule et al., 2019). This phenomenon, known as representational drift, raises a central unresolved question in image recognition. Does drift induce arbitrary changes in image representations calling for intricate downstream adaptive mechanisms, or is it structured in ways that remain compatible with stable recognition of images?

Chronic neural recordings have shown that representational drift occurs across several animal brain systems, including visual, olfactory, parietal, and hippocampal circuits (Bauer et al., 2024; Berners-Lee et al., 2023; Climer et al., 2025; Driscoll et al., 2017; Marks and Goard, 2021; Schoonover et al., 2021; Zheng et al., 2026). In the visual system, mouse studies have shown that neural responses to naturalistic movies drift across sessions (Aitken et al., 2022; Deitch et al., 2021; Xia et al., 2021), despite relative stability for simpler stimuli such as gratings (Bauer et al., 2024; Marks and Goard, 2021). During naturalistic movie watching, both calcium imaging and neuropixels recordings reveal drift from timescales of seconds to days (Deitch et al., 2021). Evidence in humans remains more limited although one recent longitudinal functional magnetic resonance imaging (fMRI) study focusing on low-level visual information showed that natural-scene representations in primary visual cortex drift over months (Roth and Merriam, 2023). EEG provides a complementary window into representational drift because it measures temporally resolved responses noninvasively and is widely used to study cognitive mechanisms in humans (Cichy and Pantazis, 2017). Therefore, characterizing drift using this modality could have implications for a broad range of cognitive neuroscience studies in humans. Moreover, image-evoked EEG responses reflect the millisecond-scale time-course of visual processing, providing a temporal dimension that is not accessible with fMRI. Whether image-evoked representations measured with EEG exhibit analogous drift across sessions remains unknown.

Moreover, the functional consequences of drift are known to depend on its pattern. Drift that is independent across response channels could substantially reorganize image representations, whereas coordinated drift may preserve global features of the representation, a distinction that has implications for downstream readout. Prior work suggests that drift can be constrained in ways that preserve decodable information, but fixed decoders may still degrade without adaptive downstream mechanisms (Gallego et al., 2020; Micou and O’Leary, 2026; Qin et al., 2023; Rule et al., 2020; Rule and O’Leary, 2022; Zaid and Schaffer, 2026). Whether image-evoked EEG drift is coordinated across channels, which properties of the representational space of images, if any, are spared by drift, and the constraints they impose on downstream compensatory mechanisms needed to maintain stable percept remain unknown.

Large-scale datasets such as THINGS-EEG2 now make it possible to ask whether repeated image presentations evoke stable or drifting multichannel response patterns across recording sessions (Gifford et al., 2022). Here, we analyzed image-evoked EEG response patterns and found that image representations drifted across sessions, that drift was structured and aligned with image-coding dimensions, and that it impaired the ability of a fixed decoder to generalize across sessions. However, drift preserved the statistics of pairwise distances between images and the dimensionality of image representation. Using a mechanistic model of a downstream network that learns via Hebbian/anti-Hebbian plasticity, we further show that such structured drift allows for faster adaptive compensation. Our findings suggest that although drift in image representations is aligned with image-coding dimensions, its structure makes it easier for downstream circuits to compensate, thereby facilitating stable recognition.

## 3 Results

### 3.1 Decoding image identity from EEG responses

We analyzed the THINGS-EEG2 dataset (Gifford et al., 2022), in which ten participants viewed *N* = 200 images across four EEG recording sessions. Recordings were made at 1000 Hz and downsampled to 100 Hz; we analyzed responses from *K* = 17 scalp electrodes overlying occipital and parietal cortex (Figure 1A-C). Each session included 20 repetitions of each image, with images presented for 100 ms followed by a 100 ms gap during which the screen was blank (Methods). We first asked whether and when image identity was most strongly represented in *K*−dimensional multi-channel response patterns by using linear support vector machines (SVM) to perform cross-temporal decoding for each image pair (Figure 1D). Decoding performance rose significantly above chance within 75.0±1.7 ms after image onset, indicating that the multi-channel response patterns carried reliable image-specific information (Figure 1E). However, this information was temporally localized. Decoders trained at one time point generalized poorly to other time points with a half-decay timescale of 27.5±1.2 ms (Figure 1F), suggesting that EEG image representations were quite dynamic. Across all image pairs and participants, the cross-validated decoding performance peaked to an accuracy of 69.6 ± 1.1% at 114.0±1.6 ms after image onset suggesting that responses carried substantial information about image identity at that time point (Figure 1G). All subsequent analyses were therefore performed using responses at this time point unless specified otherwise.

**Figure 1:**
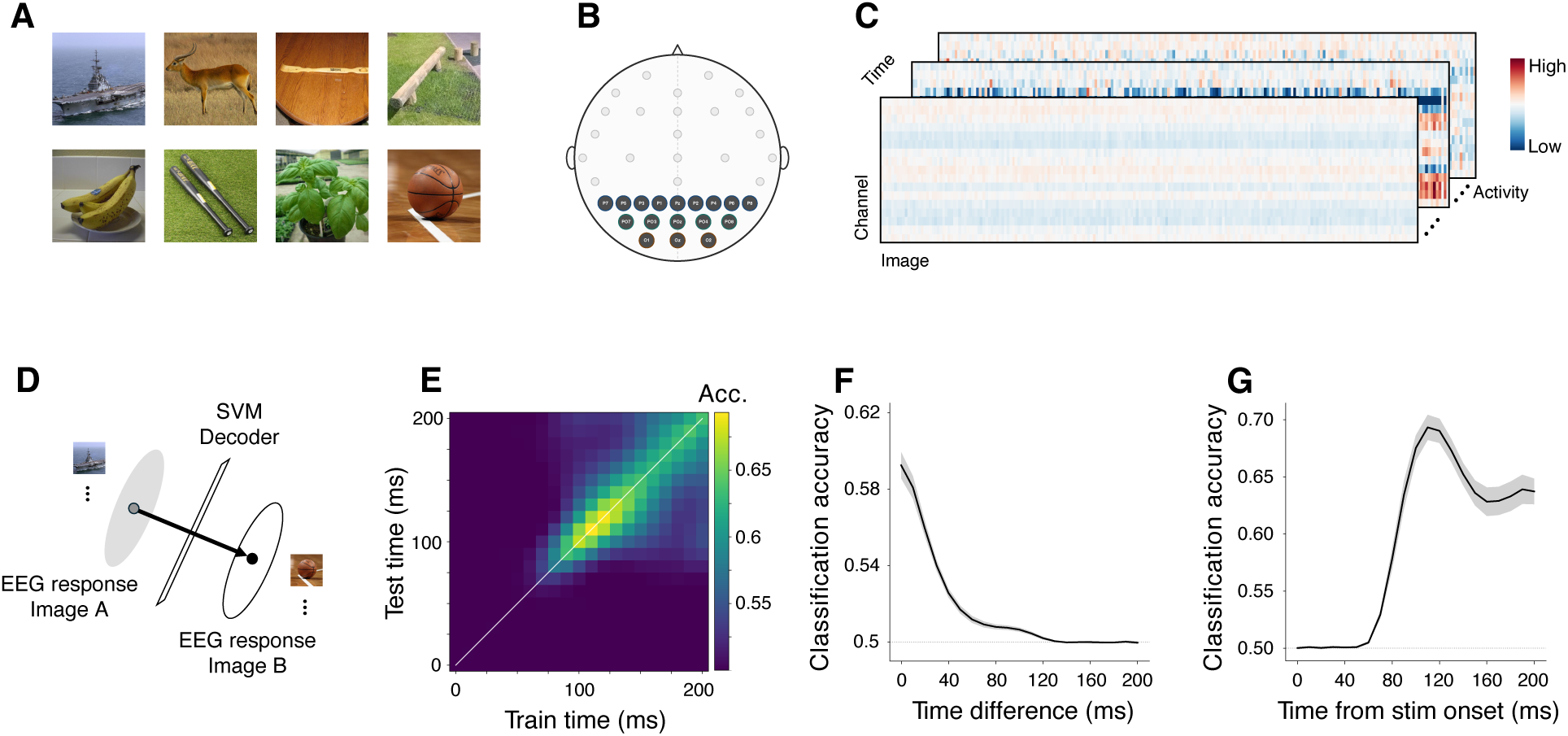
Cross-session EEG recording on visual object recognition task and EEG decoding performance. We used a publicly available dataset called THINGS-EEG2, with EEG responses to same images across four sessions. (**A**) Example stimuli from THINGS-EEG2: the dataset contains 200 images spanning 27 categories. (**B**) The 17 EEG channels used for analysis, located over parieto-occipital cortex. (**C**) Data from a single session is a 3-D array (channels × images × time): for each image, the 17-channel response was sampled every 10 ms from stimulus onset (0 ms) to 200 ms. To investigate whether it is feasible to decode image identity from EEG responses, we constructed a within-session image-wise decoder across all time points from onset of the image presentation to 200 ms after. (**D**) Linear SVM decoders were trained to perform pairwise classification between responses to individual pairs of images at each time point (0-200ms). (**E**) A heatmap of cross-temporal decoding accuracy averaged across image pairs, showing that the decoding accuracy rises about ≈ 70 ms after image onset, with limited generalization to nearby time points. (**F**) Cross-validated decoding performance was temporally localized with a decay time-constant of around 25 ms. (**G**) Classification accuracy across all pairs of images showed that the peak accuracy was around 110 ms and it was consistent across participants (Supplemental Figure S1).

### 3.2 Coordinated representational drift in EEG responses

Previous studies of neurons in mouse visual cortex have demonstrated that representations of naturalistic movies drift over time (Aitken et al., 2022; Marks and Goard, 2021; Xia et al., 2021). A recent human fMRI study of primary visual cortex likewise reported representational drift of naturalistic scenes (Roth and Merriam, 2023). We therefore hypothesized that image representations in EEG responses identified above also drifted across recording sessions. For each image, we computed a trial-averaged multichannel *K*−dimensional response vector by averaging responses across trials within a given session. We then quantified cross-session representational stability of that image by estimating the cosine similarity between response vectors from pairs of sessions. To control for random variability across trials within session, we also estimated within-session representational stability by estimating the cosine similarity between the mean response vectors from non-overlapping subsets of trials from the same session. We found that similarity between cross-session response patterns was significantly lower than that of within-session response patterns (within-session: 0.51±0.03, cross-session: 0.41±0.02, mean drop in similarity: 0.1±0.02, *p* = 0.002, two-sided paired permutation test (*N* = 10); Figure 2A). This reduction in cross-session representational stability of individual images indicates that image-specific EEG response patterns drifted across sessions, and this drift cannot be attributed to random variability across trials within a session.

**Figure 2:**
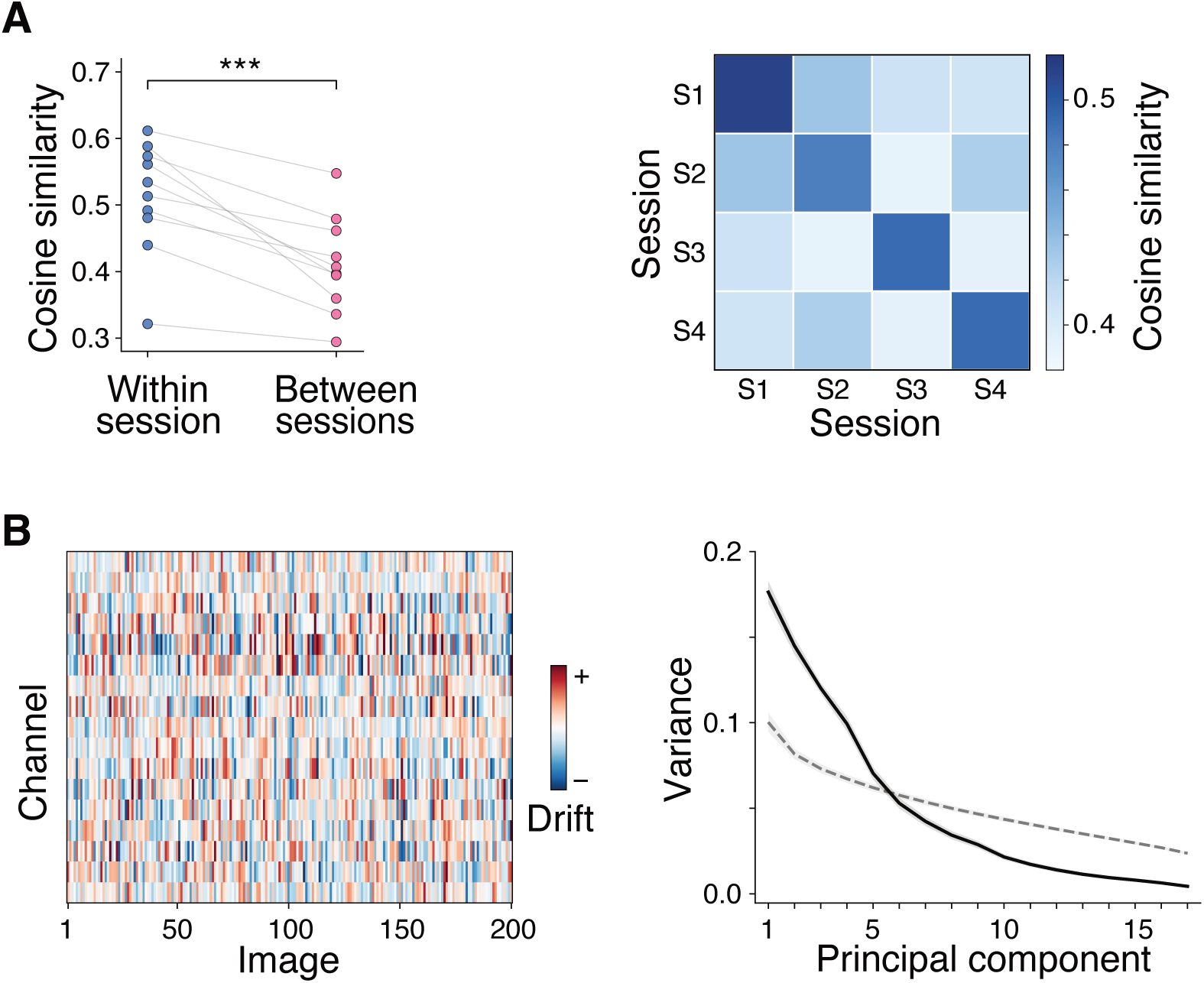
Coordinated drift in cross-session EEG recordings. (**A**) Left: Comparison of the cosine similarity between image representations computed within session (blue) or across pairs of sessions (pink). Each data point corresponds to one participant. Right: Cosine similarity for each pair of sessions, averaged across participants. (**B**) Left: The drift matrix computed using an example pair of sessions from one participant. Right: Fraction of variance explained as a function of principal components (solid line denotes data and dashed line denotes channel-shuffled control). Results were consistent across participants (Supplemental Figure S2).

Studies of neural populations have shown that representational drift can be coordinated across neurons, rather than reflecting independent changes to response properties in each neuron (Chambers and Rumpel, 2017; Eppler et al., 2025; Keinath et al., 2022; Qin et al., 2023; Rule et al., 2020). We next asked whether analogous coordination was evident at the level of EEG channels. For each image and each pair of sessions, we defined a *K*−dimensional drift vector as the difference between the mean multi-channel response vectors from the two sessions (Methods). We then constructed a *K* × *N* drift matrix containing these response vectors across all *N* images (Figure 2B – left). If drift were uncoordinated across channels, many dimensions would be needed to explain the variance in this matrix. In contrast, coordinated drift would be expected to concentrate variance along a smaller number of dominant modes. Applying principal component analysis to the drift matrix revealed that the top 5 principal components (PC) alone explained 71% of the variance, suggesting that cross-session changes in image representations were structured and coordinated across EEG channels (Figure 2B – right). Across subjects, the participation ratio (PR) of the eigenvalue spectrum was significantly lower than that of the channel-shuffled control (data: 8.3 ± 0.9, shuffled: 14.6 ± 0.5, *p* = 0.002, two-sided permutation test (*N* = 10)). A stronger concentration of variance in the top PCs compared to channel-shuffled controls suggests that the cross-session drift was coordinated across EEG channels.

### 3.3 Drift aligns with stimulus-coding dimensions and impairs decoding

Since drift was coordinated, we asked whether it was predominantly aligned with response dimensions that encode image identity. To identify these image-coding dimensions, we first applied principal component analysis to the *K* × *N* response matrix, where rows corresponded to multi-channel response patterns and columns to images. We then projected each image-specific drift vector onto the resulting stimulus principal components to quantify the fraction of total drift variance explained by each component (Methods). If drift were unrelated to image-coding dimensions, then drift vectors would show similar alignment to all the components (Figure 3A – dashed). In contrast, if drift was concentrated within image-coding dimensions, drift vectors should preferentially align with the leading principal components. We found that drift was disproportionately concentrated in the subspace spanned by the leading stimulus principal components. Each of the top six principal components accounted for a larger fraction of drift than expected by chance, indicating that drift was not distributed uniformly across the EEG response space (Figure 3A – solid). Instead, it was predominantly concentrated in the low-dimensional response subspace that encoded image identity.

**Figure 3:**
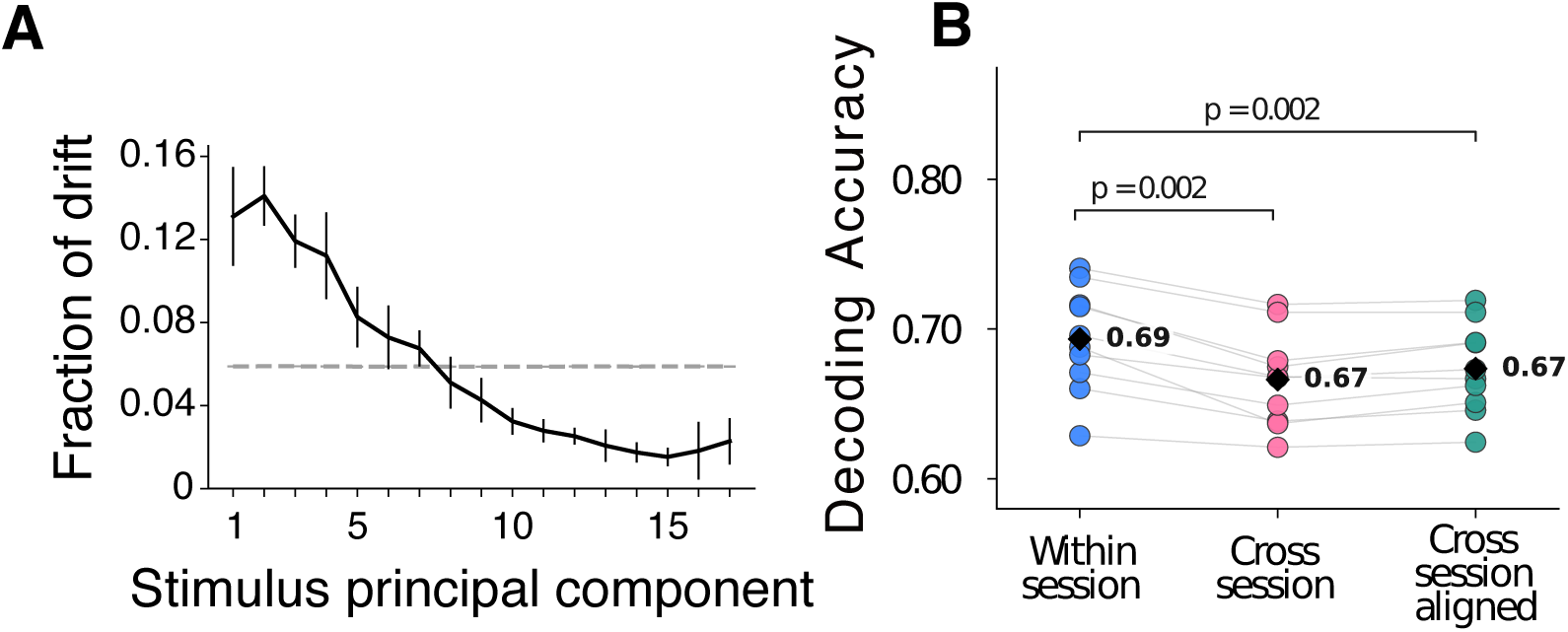
Drift is aligned with the stimulus subspace and decreases decoding accuracy. Gray dashed line denotes alignment expected by chance. (**A**) The alignment of the drift vector with the different stimulus principal components. (**B**) Decoding accuracy of classifiers trained and tested within the same session (blue), across sessions (pink), or across session after applying a global linear transformation to align the response matrix from the pair of sessions (green).

In theory, drift that is aligned with stimulus-coding dimensions should be detrimental to decoding stimulus identity (Rule et al., 2020). Concretely, if image-specific EEG responses change across sessions along dimensions that also distinguish image identity, then a decoder trained in one session may no longer generalize to later sessions. To test this, we performed pairwise image decoding by training a linear SVM and testing its performance either within the same session or across different sessions. As predicted, classifiers trained and tested within the same session achieved significantly higher decoding accuracy than classifiers trained in one session and tested on another session (mean within-session accuracy: 0.69, cross-session: 0.67, mean drop in accuracy: 2.7 ± 0.4 %, *p* = 0.002, two-sided paired permutation test (*N* = 10); Figure 3B - blue vs pink). This suggests that drift compromises image representation and impairs decoding. To test whether the decoding discrepancy between within and cross session could potentially arise from the movement of the electrodes across the scalp across sessions, we applied a global linear transformation to align cross-session response matrices. Concretely, we recomputed the cross-session decoding accuracy for each pair of sessions after aligning the response matrix from the later session to that of the previous session (Methods). The mean cross-session decoding performance after alignment remained significantly lower than within-session decoding performance (mean drop in accuracy: 2.0 ± 0.3 %, *p* = 0.002, two-sided paired permutation test (*N* = 10); Figure 3B - blue vs green), suggesting that cross-session degradation in decoding cannot be attributed to global movement of EEG electrode array.

### 3.4 Drift preserves the dimensionality of image representations

Although representational drift was aligned with stimulus-coding dimensions and impaired decoding, the average loss in decoding performance across images was quite modest with a mean drop in accuracy of less than 3%. To obtain a granular understanding of the effects of drift, we examined the direction of image-specific drift relative to the decoding axes. For each image, we measured the cosine similarity between its drift vector across a pair of sessions and the SVM decoding axes, learned in the original session, that discriminated that image from every other image (Methods). Decoder axes were oriented so that positive alignment indicated movement toward the mean response of the other image. In other words, positive cosine similarity corresponds to drift that reduces the separation between image pairs and impairs decoding, whereas negative cosine similarity corresponds to drift that increases the separation (Figure 4A). We found that image pairs with more positive alignment between the direction of drift vector and decoding axis and larger drift magnitude, showed a greater reduction in cross-session decoding accuracy (*r*_direction_ = −0.47*, p <* 0.0001; *r*_magnitude_ = −0.26, *p <* 0.0001; Wald test). Notably, the impact of drift was heterogeneous across image pairs. Decoding performance improved for 34 ± 7% of pairs, decreased for 57 ± 8%, and remained unchanged for 8 ± 1% (Figure 4B). Therefore, modest drop in average decoding performance was due to the heterogeneous effects of drift rather than weak drift. The heterogeneity further suggests that the pattern of drift might be constrained by the geometry of the representational space. We therefore next asked whether global geometric features such as pairwise distances and dimensionality were preserved across sessions.

**Figure 4:**
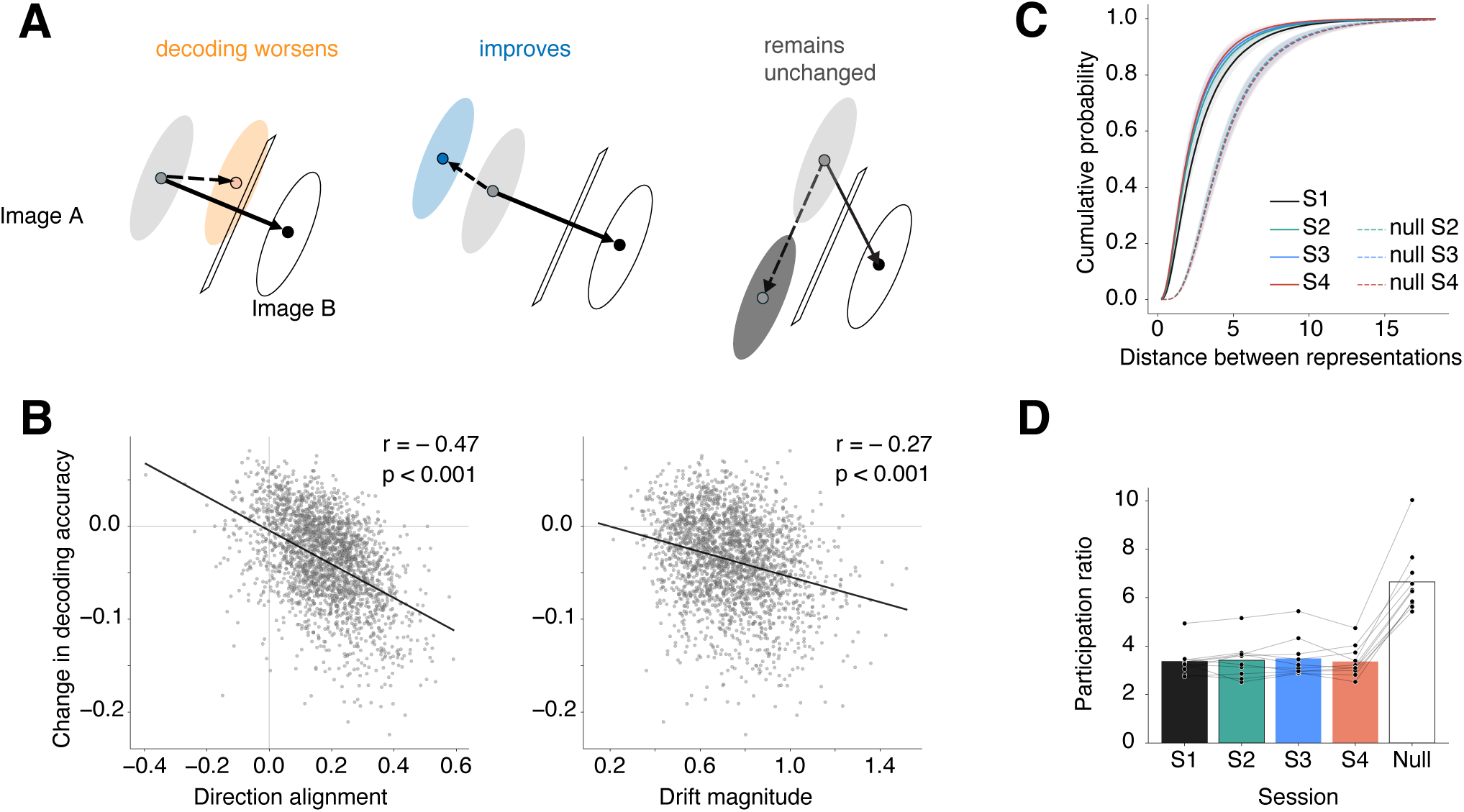
Drift is heterogeneous but preserves dimensionality. (**A**) Possible ways drift might influence decoding: drift aligned with the decoding axis impairs decoding accuracy; anti-aligned drift increased the margin improving decoding accuracy; drift along the orthogonal direction leaves decoding performance unchanged. (**B**) Change in decoding accuracy as a function of the alignment between drift vector and decoding axis (left) or as a function of drift magnitude (right). Direction alignment was defined as the cosine similarity between drift vector and decoding axis; Drift magnitude was defined as the norm of the drift vector. (**C**) The distribution of pairwise distances between image representations for various sessions of data and simulated null responses (see text). (**D**) Dimensionality (participation ratio) of image representation for various sessions of data and simulated null responses.

To test this, we computed an *N* × *N* representational dissimilarity matrix (RDM) for each session, where each entry corresponded to the distance between mean response vectors to an image pair. As a null model, we generated pseudo-RDMs using simulated drift with the same image-specific drift magnitudes as the data, but with drift directions shuffled across images (Methods). We found that the distribution of pairwise distances between image representations was stable across sessions but shifted substantially under the null model of drift. Mean Wasserstein distances (relative to session 1) were 0.61 ± 0.13, (95%CI ∈ [0.33, 0.90]) for raw data and 1.77 ± 0.06, (95%CI ∈ [1.62, 1.91]) for shuffled data, with a significant difference between mean distances of −1.15 ± 0.11 (*p* = 0.00195, two-sided paired permutation test (*N* = 10); Figure 4C). Therefore, null responses simulated by matching drift magnitude but randomizing drift direction substantially increased the average pairwise distances between image representations. This indicates that the volume of the image-coding subspace might be preserved across sessions in spite of representational drift. To directly test this, we asked whether the dimensionality of image representations was stable across sessions.

We quantified dimensionality using the participation ratio of the image response matrix in each session. Empirical responses had a lower participation ratio than randomized-direction null responses (3.4 ± 0.2 versus 6.7 ± 0.4, mean ± SEM across participants; two-sided paired permutation test, *p* = 0.0015). The empirical participation ratio did not differ significantly across sessions (*p* = 0.78, Friedman repeated-measures test; Figure 4D), indicating that representational drift preserved the dimensionality of image-evoked EEG response patterns.

### 3.5 Structured drift allows for faster adaptive compensation

Because drift preserved the dimensionality of image representations, we asked whether this structure might confer a performance advantage over unstructured drift. Decoding performance could, in principle, be preserved trivially if drift occurred mostly orthogonal to image-coding dimensions, in which case it would have little impact on decoding. However, this was not the case in the data. We showed earlier that drift was concentrated along image-coding dimensions impairing the ability of decoders to generalize across sessions. We therefore hypothesized that the structure of drift may instead facilitate adaptive compensation by unsupervised downstream mechanisms, allowing decoding performance to recover more rapidly than they would under unstructured drift.

Hebbian/anti-Hebbian networks can extract principal components from their inputs, and previous theoretical studies have proposed such unsupervised learning rules as a potential mechanism by which downstream areas transform drifting representations into stable ones (Rule et al., 2020). While it remains to be seen whether the structure of drifting EEG image representations observed here also applies at the neuronal level, we used this framework to test whether structured drift could be more amenable to downstream compensation. We constructed a two-layer neural network model in which input units were subject to representational drift, while units in a recurrent hidden layer underwent Hebbian/anti-Hebbian plasticity and provided input to a fixed linear decoder (Methods; Figure 5A) (Leen, 1991). The network consisted of feedforward connections governed by Hebbian plasticity and recurrent lateral connections governed by anti-Hebbian plasticity. Hebbian learning allowed the feedforward weights to extract principal directions of the input representation, whereas anti-Hebbian plasticity decorrelated the activity of output neurons and prevented multiple neurons from converging onto the same leading principal component. We evaluated the model’s ability to compensate for representational drift that was either structured or unstructured. Because our goal was to test the implications of drift structure rather than the mechanisms that generate it, we modeled drift as an abrupt perturbation of the input representation rather than as a slowly evolving process. In the structured condition, drift preserved the dimensionality of the input representation by maintaining the shape of the covariance eigenspectrum while allowing the eigenvectors to change (Figure 5B, participation ratio (PR) of 2.5). In the unstructured condition, drift increased the dimensionality of the input representation (Figure 5B, PR of 5.0). Structured and unstructured drift initially reduced decoding performance to a comparable extent (Figure 5C). Strikingly, however, performance recovered to its original level substantially faster after structured drift (Hebbian/anti-Hebbian model: *τ*_unstructured_*/τ*_structured_ = 5.09 ± 0.33, *p <* 0.0001, two-sided paired permutation test, Figure 5C). We found qualitatively similar results in a model in which downstream compensation was implemented using the generalized Hebbian learning algorithm (Sanger, 1989)(*τ*_unstructured_*/τ*_structured_ = 1.66±0.03, Supplementary Figure S4). These results suggest that preserving the dimensionality of image representations can facilitate more rapid adaptive compensation by downstream unsupervised mechanisms.

**Figure 5:**
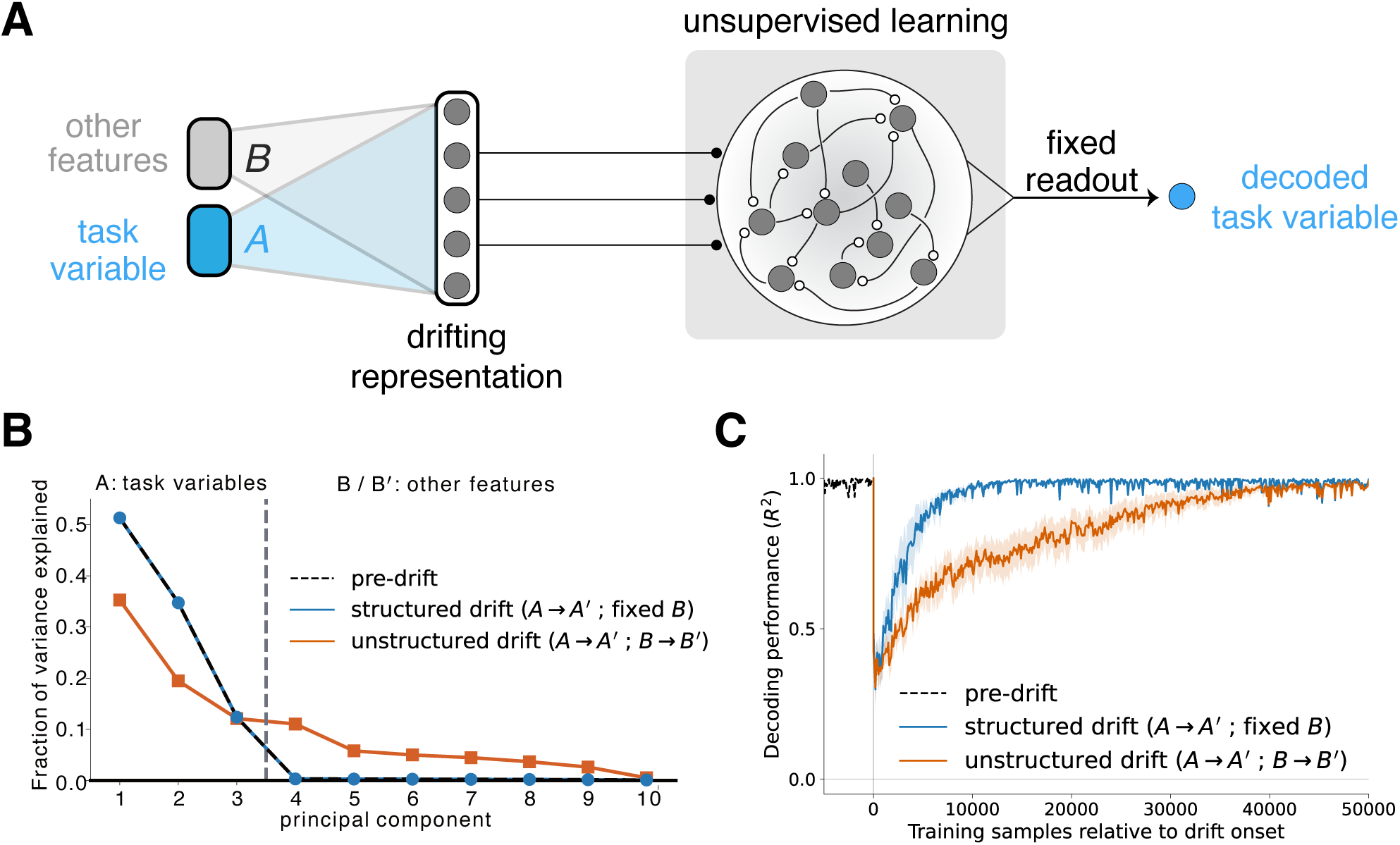
Adaptive compensation by unsupervised learning depends on the structure of drift. (**A**) Model schematic. A stimulus comprising a task variable (embedded by *A*) and additional task-irrelevant features (embedded by *B*) is projected onto a neural population whose representation drifts over time (Methods). A Hebbian/anti-Hebbian network was trained on the population activity both before and after drift onset, and a fixed linear readout decodes the task variable from its output. Because the readout is held fixed, sustaining good decoding performance requires the circuit to compensate for drift by extracting stable, task-relevant features from the drifting representation. (**B**) Eigenvalue spectrum of the population representation under structured and unstructured drift. For structured drift, we held the other features *B* fixed during drift generation and applied only a rotation to the covariance matrix (*A* → *A*^′^), preserving the eigenvalue spectrum of the pre-drift condition (blue overlaps the dotted line). For unstructured drift, we additionally let *B* diffuse (*B* → *B*^′^), causing the dimensionality to expand (orange). (**C**) Decoding performance (*R*^2^) of the fixed readout as a function of the number of training samples presented to the circuit, aligned to drift onset. Both forms of drift produce an abrupt drop in decoding accuracy from the pre-drift baseline (dotted) at onset, followed by recovery as the circuit realigns to the new representation. Recovery is faster under structured drift (blue) than under unstructured drift (orange). Solid lines show the mean across 50 simulations and shaded regions shows ±1SEM.

## 4 Discussion

Our finding of representational drift in human EEG responses complements a growing body of work showing that neural representations change over time despite stable sensory input and behavior. In the mouse visual system, chronic neural recordings have shown that stimulus-evoked responses in visual cortex drift across days (Bauer et al., 2024; Marks and Goard, 2021). Notably, such drift is more apparent for naturalistic stimuli than for classic orientation gratings (Marks and Goard, 2021), suggesting that representations of complex visual input may be particularly susceptible to change.

Recent longitudinal studies provide growing evidence that human neural representations also vary across recording sessions. An fMRI study showed that representations of natural scenes in primary visual cortex drift gradually over months (Roth and Merriam, 2023), providing important evidence for long-term changes in human visual representation. A limitation of that study is that individual images were sparsely repeated across sessions. Drift was therefore inferred from changes in tuning to low-level features derived from image-computable encoding models rather than from repeated measurements of the same image-evoked response. Moreover, the temporal resolution of fMRI makes it difficult to distinguish changes in early stimulus-evoked activity from changes emerging through later recurrent or feedback processing (Cichy et al., 2014). In the dataset analyzed here, each image was presented 80 times across four sessions, allowing us to detect the timepoint at which images were maximally discriminable and quantify session-to-session drift of image-evoked response patterns at millisecond time scales. Thus, our results extend previous evidence for human representational drift by demonstrating image-specific drift in EEG responses over time. Complementary MEG work found that visually evoked response patterns varied across scanning days spanning months although they did not test whether image-level decoders were stable across days (Paulsen et al., 2026). Beyond perceptual representations, a recent fMRI study reported cross-session drift in multivariate activity patterns encoding visual working-memory load across six sessions (Adam et al., 2025). Together, these studies suggest that cross-session representational variability is a general feature of human neural activity.

These findings pose a challenge for understanding brain function: how can neural representations change across sessions while perception and behavior remain stable? One explanation for why representational drift can coexist with stable behavior is that drift occurs in dimensions orthogonal to those that encode task-relevant variables (Keinath et al., 2022; Paulsen et al., 2026; Rule et al., 2019). However, work in mouse posterior parietal cortex shows that drift in task-related variables such as position is not strictly confined to such coding-null dimensions (Driscoll et al., 2017; Rule et al., 2020). Whether sensory representations obey a similar constraint has remained unclear. Our results show that image-evoked EEG drift was substantially aligned with image-coding axes and impaired fixed-decoder generalization across sessions. Thus, stable image recognition cannot be explained solely by the idea that representational drift is orthogonal to the representational space of images. Although drift was aligned with the image-coding dimensions, its effect on decoding was modest on average because its effects were heterogeneous across image pairs. For some pairs, drift moved image representations closer together and reduced decoding accuracy while for others, drift increased their separation and improved decoding. This heterogeneity may reflect an important distinction between drift in representations of categorical and continuous variables. For continuous variables such as position, changes along the coding dimension may produce systematic errors in the represented value. In contrast, for categorical stimuli such as images, there is no single ordered axis along which all errors accumulate. Drift can therefore reduce the margin between some category boundaries while increasing others, producing a weaker average effect on decoding despite substantial movement of individual representations. This suggests that the functional consequences of representational drift may depend on the structure of the variable being represented, with categorical and continuous stimulus spaces imposing different constraints on how drift affects downstream readout. Evidence for drift in other categorical variables like olfactory representations (Berners-Lee et al., 2023; Chen and Padmanabhan, 2026; Schoonover et al., 2021; Zheng et al., 2026) provides a useful setting to test this idea further, by asking whether odor drift also produces heterogeneous changes in pairwise decoding.

A major finding of our study is that image-evoked EEG drift was structured rather than reflecting independent changes across channels. This is consistent with previous work showing that representational drift can be coordinated across neural populations and constrained by the geometry of the response space (Aitken et al., 2022; Kriegeskorte and Wei, 2021; Peters et al., 2026; Zaid and Schaffer, 2026). For example, studies in mouse visual cortex have shown that even as individual neural responses drift, the relationships between population activity patterns can remain stable (Deitch et al., 2021; Xia et al., 2021). Human fMRI work has similarly found drift in voxel-wise visual responses despite preserving population-level dissimilarities between stimuli (Roth and Merriam, 2023). Our results extend this to image-evoked EEG responses. Drift did not alter the overall distribution of pairwise distances between image representations suggesting that drift affects individual image responses while preserving the density of the representational space. One possible explanation for this preserved distance is that drift is constrained by the dimensionality of image representations. Consistent with this idea, we found that the effective dimensionality of image-evoked response patterns was stable across sessions. Preservation of dimensionality may reflect a biological constraint on the coding resources available for encoding complex visual stimuli (Gallego et al., 2020). We asked whether such structure might be useful for downstream systems that must adapt to drifting inputs. Previous theoretical work has shown that Hebbian and anti-Hebbian plasticity rules can extract and track dominant variance dimensions in streaming inputs (Becker and Plumbley, 1996; Földíak, 1989; Fyfe, 2000; Leen, 1991; Rubner and Tavan, 1989; Sanger, 1989), and related models have proposed that local plasticity can allow downstream circuits to maintain stable readout of drifting neural codes (Rule et al., 2020; Rule and O’Leary, 2022). Motivated by this work, we developed a mechanistic model in which a downstream network compensated for drift through Hebbian/anti-Hebbian plasticity. The model recovered decoding performance more rapidly when drift preserved the dimensionality of the input representation than when drift increased dimensionality. These results suggest that the specific structure of drift in the data allows for downstream circuits to more easily track drifting representations. More broadly, this means that the impact of drift on the dimensionality of representation should be a key indicator of long-term behavioral stability, a prediction that could be tested by combining longitudinal recordings with a behavioral task in humans or animals.

Our findings carry some important caveats that present opportunities for future work. A general problem in studying representational drift with noninvasive methods such as EEG and fMRI is that apparent changes in response patterns may partly reflect measurement-related factors, such as differences in electrode positioning, impedance or cross-session registration. If drift were driven entirely by such factors, one might expect it to act primarily as a global transformation of the response, such as a rotation, scaling, or other shared linear mapping across images, rather than producing image-specific changes in representational stability and decodability. Consistent with this distinction, we found that global linear transformations did not fully account for the observed pattern of drift (Figure 3B). However, a stronger signature of drift would be gradual accumulation across time, such that more widely separated sessions show larger changes, as reported in a previous fMRI study (Roth and Merriam, 2023). This was difficult to assess conclusively here because the dataset contained only four recording sessions. A related caveat is that drift in EEG response patterns need not reflect drift at the level of single neurons, because each EEG channel reflects a mixture of activity from distributed cortical sources and can be affected by changes in source orientation and projection to the scalp. Nevertheless, drift at this macroscopic scale is still important to characterize as it is central to interpreting EEG studies that characterize human perceptual and cognitive representations. Future studies with more longitudinal sessions, digitized electrode positions, source-localized EEG or MEG, and behavioral measures of recognition could help distinguish measurement drift from neural representational drift and test whether the structured changes observed here predict perceptual stability across time.

## 5 Methods

### 5.1 Dataset and EEG response vectors

We analyzed a publicly available THINGS-EEG2 dataset, which contains cross-session human EEG recordings collected during repeated presentations of natural images (Gifford et al., 2022). We used the test image set, consisting of 200 images presented repeatedly across four recording sessions in 10 participants. Stimuli were drawn from the THINGS image database and resized to 500 × 500 pixels. The test set contained natural images of recognizable object concepts, such as antelope and cupcake, spanning 27 higher-level categories, including animals and foods. Because the released preprocessed test-set files did not preserve the session ordering required for our cross-session analyses, we reconstructed the original four-session structure using publicly available preprocessing code (https://github.com/gifale95/eeg_encoding) from THINGS-EEG2 before performing the analyses reported here. EEG responses were analyzed from 17 occipital and parietal channels. Each image was presented 20 times per session, yielding 80 trials per image for each participant across the four sessions. The experiment used a rapid serial visual presentation (RSVP) paradigm. Participants viewed sequences of 20 centrally presented images while seated in front of a computer screen. Each sequence began with a 750 ms blank screen containing a central fixation point. Each image was presented for 100 ms and followed by a 100 ms blank interval before the next image appeared. Further details of the experimental design are provided in (Gifford et al., 2022). For the present analyses, we performed temporal analyses for every 10 ms time point within 0–200 ms post-stimulus window, corresponding to the full interval between the onset of one image and the onset of the next. At each time point *t*, the response to repetition *r* of image *i* in session *s*, x^s^_i,r_(*t*) contained a vector-valued activity from *K* = 17 occipital and parietal channels. The average response was given by

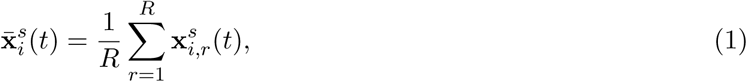

where *R*=20 denotes the number of repeated trials.

For analyses based on the full set of images, we arranged the mean response vectors into a channel-by-image response matrix **M***^s^*(*t*) ∈ R*^K^*^×*N*^ with columns populated by x^s^_i_(*t*) where *N* = 200 is the number of images. Unless otherwise stated, all analyses were performed separately for each participant and time point.

### 5.2 Quantifying drift

To estimate within-session similarity, we randomly split *R* = 20 trials for each image within a session into two non-overlapping sets of 10 trials. We averaged the responses within each split and computed the cosine similarity between the two split-half response vectors. For cross-session similarity, we sampled 10 trials each from two different sessions (*s* and *s*^′^) for the same image, averaged the responses within each session, and computed the cosine similarity between the two session-specific response estimates x^s^_i_ and x^s’^_i_. We repeated this random resampling procedure 500 times and averaged the resulting similarity estimates to reduce dependence on any particular trial split. This procedure was repeated across random resamplings, images, session pairs, participants, and time points.

To determine whether cross-session drift was coordinated across EEG channels, we constructed drift matrices across each pair of sessions *s* and *s’* as **D*^s^*** = **M*^s’^*** − **M*^s^*** where each column of **D*^s^*** represents the cross-session drift vector for one image in the *K*-dimensional EEG response space. We then applied PCA on **D***^s^* and quantified the fraction of variance fraction explained by each of the *K* principal components (PC). To construct a null distribution, we independently shuffled the image identities within each channel of the drift matrix before PCA. This procedure preserved the distribution of drift magnitudes of individual EEG channel while disrupting the image-specific covariance between channels. PCA was then applied and the fraction of variance explained by each of the *K* PCs was compared to that of raw (unshuffled) drift matrices.

### 5.3 Projection of drift onto stimulus space

To further investigate whether stable cross-session decoding was caused by drift being orthogonal to the stimulus space, we projected each drift vector onto the subspace spanned by the top *k* stimulus PCs, with *k* varying from 1 to 17, and measured the fraction of squared drift magnitude captured by that subspace.

We first mean-centered the response matrix **M***^s^* across images and applied singular value decomposition (SVD) to obtain an orthonormal basis for the stimulus representation. Here, we decomposed mean-centered the response matrix at the highest decoding accuracy point (*t*=110ms) as **M**^-^ *^s^* = **U***^s^* **Σ***^s^* (**V***^s^*)^⊤^ where **Σ***^s^* denotes the matrix of singular values for session *s* and the first *k* rows of U^s^_k_ Ξ {u_1_, …, u_k_} correspond to the top *k* stimulus PCs. To quantify whether drift occurred along stimulus dimensions, we projected each drift vector onto the subspace spanned by the top *k* stimulus PCs. For each image *i*, the drift vector ***δ****^s^* (*i*^th^ column of **D***^s^*) lies in the same *K*-dimensional EEG channel space as the stimulus PCs. The first *k* columns of **U***^s^* contain the leading stimulus principal-component directions and form the matrix U^s^_k_ ϵ R^K × k^. We quantified the cumulative fraction of squared drift magnitude captured by the stimulus-PC subspace as a function of the size of the subspace *k*. For each rank *k*, the subspace was composed by the first *k* stimulus PCs, where *k* was varied from 1 to *K* = 17, the dimensionality of the EEG channel space. For each image, this fraction was computed as the squared norm of the projected drift vector divided by the squared norm of the full drift vector:

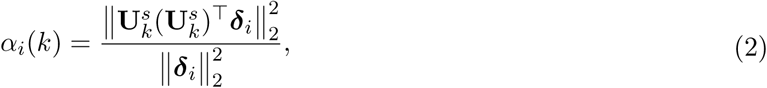

where *α_i_*(*k*) ∈ [0, 1] measures how much of the drift lies within the stimulus subspace spanned by the top *k* stimulus-coding dimensions. In addition to measuring the cumulative fraction of drift projected onto the top *k* stimulus-PC subspace, we quantified the contribution of each individual principal component. This analysis allowed us to determine whether drift was broadly distributed across many stimulus dimensions or concentrated along a small number of dominant PCs. For each image *i*, we computed the squared projection of the drift vector *δ_i_* onto the *k*-th stimulus PC direction **u***_k_*, normalized by the total squared drift magnitude as β_i_(k) = (u^T^_k_δ_i_)^2^/.‖δ_i_‖^2^_2_

### 5.4 Pairwise decoding of images

#### Within and cross-session pairwise decoding

To test whether image-specific EEG representations generalized across recording sessions, we performed pairwise image decoding using a linear support vector machine (SVM). For within-session decoding, classifiers were trained and tested on trials from the same session using stratified five-fold cross-validation. For cross-session decoding, classifiers were trained on trials from one session and tested on trials from a different session. To make within-session and cross-session decoding directly comparable, we matched the number of training and test trials across the two analyses, thereby controlling for potential biases due to differences in sample size. Decoding was performed for each pair of image, at each time point and for each participant. With 200 images, this yielded (200 _2_) = 19,900 binary classifiers per subject and target session at each time point. To summarize decoding performance at the level of individual images, we computed an image-wise decoding score by averaging the decoding accuracies across all pairwise classifiers involving that image. Because each image was paired with the remaining 199 images, each image-wise score was based on 199 pairwise decoding accuracies. For each image pair (*i, j*), we trained a linear SVM to discriminate trial-level EEG responses to the two images. The corresponding SVM weight vector, w^s^_i,j_(t), was taken as the decoding axis for that image pair in session *s* at time point *t*.

To quantify the representational drift of a given image *i* across sessions, we computed the change in the mean response vector of a given image. For example, the drift vector between sessions *s* and *s*^′^ is defined as δ^s⟶s’^_i_(t) = x^s^_i_(t). The magnitude of the drift vector for image *i* was defined as the norm of the drift vector, normalized by the average Euclidean distance between image *i* and all other image means.

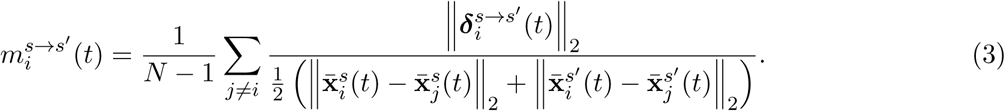

The alignment between the drift vector and the SVM decoding axis was quantified using cosine similarity. For each pair of images (*i, j*), we computed the cosine similarity between this drift vector and the SVM decoding axis w^s^_ij_(t) trained to discriminate image *i* from image *j*. To obtain an image-wise alignment score, we averaged these pairwise cosine similarities over all other images *j* ≠ *i*. Thus, each image-wise alignment value was based on *N* − 1 = 199 pairwise decoding axes, and this procedure was repeated for all *N* = 200 images.

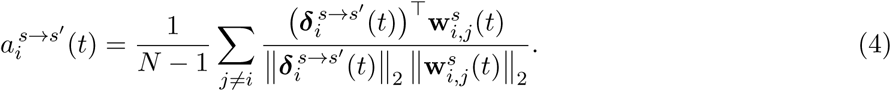

#### Global linear transformation for cross-session alignment

In longitudinal EEG recordings, differences in electrode placement across sessions could change EEG responses patterns and cause a decrease in cross-session decoding accuracy. To account for global drift caused by shifts in electrode placement, we performed an orthogonal Procrustes alignment between pairs of sessions and then tested the image-wise decoding accuracy on EEG response matrix for each participant after alignment.

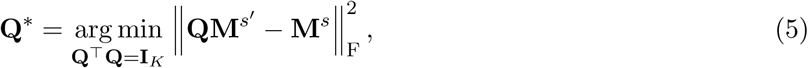

where ‖·‖_F_ denotes the Frobenius norm and **I***_K_* is the *K* × *K* identity matrix. The aligned response matrix was therefore M^s’^_aligned_ = Q*M^s’^. We then recomputed image-wise decoding accuracy by testing the classifiers trained on session *s* on the aligned response in session *s*^′^ (Figure 3B).

#### Simulation of drift vector with randomized direction

To assess how drop in decoding performance depended on the structure of representational drift, we generated simulated target-session responses with synthetic drift. The simulated responses were constructed by preserving the trial-by-trial variability from the target session (*s* ∈ {2, 3, 4}) while shifting its mean response along a random drift direction as:

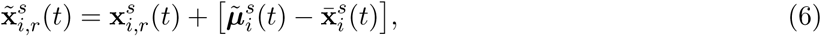

where μ^s^_i_(t)’ x^1^_i_(t) + m_i_u_i_(t) and ‖u_i_(t)‖_2_ = 1. Thus, the simulated mean response vector was generated by anchoring each image at its session 1 mean response, **x**^-1^(*t*) and adding a random unit vector u_i_(t) in *K*-dimension EEG space and a scalar drift magnitude parameter *m_i_*sampled from the empirical EEG drift magnitude distribution. For each participant, image pair, time point, and target session *s* ∈ {2, 3, 4}, we trained a linear SVM on Session 1 trials and tested it on either real or simulated target-session trials. The cross-session pairwise decoding in simulated trials was performed in the same way as the decoding schema with the raw EEG response.

### 5.5 Representational similarity and distance geometry

For each session, we constructed an representational dissimilarity matrix (RDM) containing squared Euclidean distances between mean image-response vectors. We extracted its off-diagonal lower-triangular entries and sorted them in ascending order. We quantified differences between the resulting distance distributions using the first Wasserstein distance:

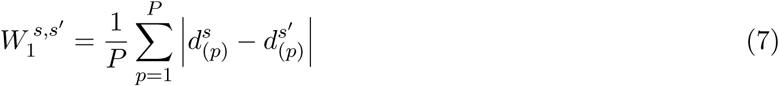

where *P* = *N* (*N* − 1)*/*2 is the number of unique image pairs, and d^s^_(p)_ denotes the *p*-th smallest squared distance in session *s*. We compared each later session with session 1 separately for each participant (Figure 4C).

### 5.6 Effective dimensionality

To quantify the dimensionality of the EEG signal subspace, the effective dimensionality was quantified as the participation ratio from the eigenvalue spectrum of the image-by-channel response matrix **M***^s^*(*t*) ∈ R*^K^*^×*N*^. The matrix was centered across images for each channel before PCA was applied. Participation ratio was calculated as:

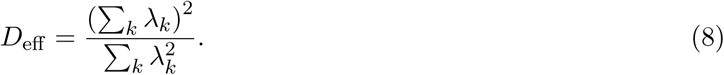

where *λ_k_* denotes the eigenvalue for component *k*. This measure estimates the number of equally weighted principal components that would produce the same variance concentration as the observed PCA spectrum. Lower values indicate that signal variance is concentrated in a small number of leading components, whereas higher values indicate that variance is distributed more evenly across components.

### 5.7 Hebbian learning simulations with empirical constraints

We simulated a Hebbian learning model to test whether the impact of representational drift on dimensionality affects the efficiency of adaptive compensation. We constructed synthetic responses whose dimensionality and variance structure were constrained by the empirical findings, and compared learning after two forms of drift: a structured drift condition that preserved the low-dimensional structure observed in the EEG data, and an unstructured drift condition that expanded the effective dimensionality of the response space.

#### Empirical constraints on the simulated data

The simulated activity was constrained by the empirically estimated covariance spectrum of EEG responses. In EEG data, image-specific response vectors were constructed by averaging repeated responses to each image within each session. PCA was then applied to these image-specific response vectors to estimate the dimensionality and eigenvalue spectrum of the empirical response space. The empirical EEG sessions showed a low-dimensional response structure, but randomized-direction null responses had approximately twice the effective dimensionality of empirical EEG responses (Figure 4D). We therefore constructed simulated representations with a comparable twofold contrast in effective dimensionality. Structured drift preserved the pre-drift covariance spectrum and effective dimensionality *D*_eff_ ≈ 2.5. Unstructured drift redistributed variance across latent components to approximately double effective dimensionality with *D*_eff_ ≈ 5, while retaining the same number of components and total variance.

#### Simulated responses

Synthetic neural activity was generated from Gaussian latent samples with a prescribed covariance. Ten latent components were linearly mixed into a population of *N* = 64 input neurons along orthonormal eigenvectors of the population covariance matrix according to **x** = **Az***_A_* + **Bz***_B_* where three components (**z***_A_*) represented task-related activity and the remaining seven (**z***_B_*) represented other features. The eigenvectors of **A** and **B** were orthonormal to each other, and the task-related and other-feature subspaces were mutually orthogonal. The simulated population had a concentrated low-dimensional task-related components accounting for more than 95% of total variance prior to applying drift.

#### Simulated drift

We applied two forms of representational drift. Structured drift was applied by rotating the eigenvectors spanning the task-relevant subspace *A* yielding **x**_str_ = **A**^′^**z***_A_* + **Bz***_B_*. Structured drift rotated the task-related eigenvectors within their three-dimensional subspace, changing **A** to **A**^′^ while keeping **B** and all latent values fixed. This transformation changed the coordinates of population activity across neurons while preserving the covariance spectrum and pairwise inner products between sample representations. This manipulation was motivated by similarity-matching models, in which orthogonally related representations can have the same objective value and noisy learning can produce coordinated drift (Qin et al., 2023). Unstructured drift applied by additionally replacing **B** with new eigenvectors **B**^′^ that remained orthogonal to the task subspace yielding **x**_unstr_ = **A z***_A_* + **B z***_B_*, where the latent variables are rescaled to **z***_A_*and **z***_B_* to obtain a covariance spectrum with higher dimensionality, *D*_eff_. Both conditions retained ten components and the same total variance. However, unstructured drift distributed more variance across the other-feature components, increasing the participation ratio from approximately 2.5 to 5. This approximately twofold increase was chosen to reflect the empirical dimensionality contrast between biological and randomized EEG drift vectors, with randomized drift exhibiting approximately twice the participation ratio of biological drift (See Figure 4D). Thus, structured drift preserved effective dimensionality, whereas unstructured drift increased it.

#### Learning rules

We compared two biologically motivated families of learning rules. In both models, input population activity was denoted by **x** and the learned output representation was denoted by **y**. First, we used a Hebbian/anti-Hebbian network with feedforward learning and lateral decorrelation Leen (1991) as shown in (Figure 5A). In this model, output activity is influenced by both feedforward input and lateral interactions among output units as **y**^(*τ*+1)^ = **Wx** + **V y**^(*τ*)^ and *V_ii_* = 0 where **V** denotes the lateral weight matrix and *τ* indexes time. The diagonal entries of **V** are fixed to zero. Feedforward weights are updated using a Hebbian learning rule with activity-dependent normalization:

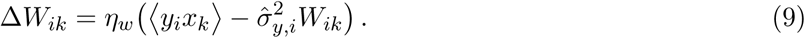

where *η_w_* is the feedforward learning rate and σ^2^ is an activity-dependent estimate of the variance of output unit. Lateral weights were updated using an anti-Hebbian decorrelation rule:

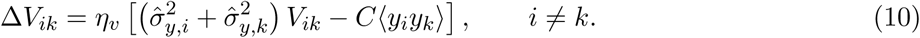

where *η_v_* is the lateral learning rate and *C* controls the strength of decorrelation. In addition to the above local, online Hebbian/anti-Hebbian mechanism, we used Sanger’s rule, an online Hebbian learning rule that extracts ordered principal components from the input activity (Sanger, 1989), see Figure S4. In this case, the output activity was given by **y** = **Wx** where **W** denotes the feedforward weight matrix. The feedforward weights were updated according to:

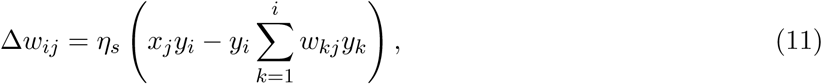

where *η_s_* is the learning rate. This rule allows successive output units to learn ordered, decorrelated components of the input covariance structure.

#### Task variable and downstream readout

To evaluate whether the simulated population supported stable downstream decoding, we defined a scalar task variable from the original latent representation **z**. The same task variable was used across both drift conditions, so changes in decoding performance reflected changes in the neural representation rather than changes in the task itself. After the initial learning phase, a linear downstream decoder was trained to predict the task variable from the learned output representation **y**. This decoder was then held fixed during the post-drift phase. The Hebbian models were allowed to update responses **y** after drift, but the downstream readout was not retrained. This design tested whether local adaptive learning could restore a task-relevant representation after representational drift.

#### Training procedure

Each simulation consisted of an initial learning phase followed by a post-drift adaptation phase. During the initial phase, each model was trained on samples drawn from the original low-dimensional input distribution. After this phase, the downstream decoder was fit and then frozen. Drift was then introduced by replacing the input distribution with either the real-like low-dimensional drift distribution or the random-like expanded-dimensionality drift distribution. The Hebbian models continued to update their internal weights in response to the drifted input activity. For the Hebbian/anti-Hebbian model (Leen, 1991), both feedforward and lateral weights were updated. For Sanger’s rule (Sanger, 1989), only the feedforward weights were updated. Model performance was evaluated throughout the post-drift phase.

#### Evaluation of adaptive compensation

Model performance was quantified by the coefficient of determination, *R*^2^, between the true task variable and the decoded task variable. For each learning model and drift condition, we measured the trajectory of *R*^2^ during post-drift adaptation. Learning efficiency was defined as the number of training samples or update steps required to recover to a fixed fraction of the pre-drift decoding performance. We compared learning efficiency between the real-like and random-like drift conditions. This comparison tested whether dimensionality expansion increases the sample complexity of adaptive compensation, even when the immediate impairment of fixed decoding is comparable between real and randomized drift.

### 5.8 Statistical analysis

EEG analyses were first carried out within each subject, and group-level inference was performed across subjects. Statistical results were reported as p-value from two-sided permutation test unless stated otherwise, with threshold for statistical significance set to 0.05.

For the within-session versus cross-session cosine similarity analysis, subject-level estimates were obtained using repeated split-trial resampling. Within-session reliability was estimated by randomly dividing the 20 repeated trials from a given session into two non-overlapping sets of 10 trials, averaging the response within each split, and computing the cosine similarity between the two split-half response vectors. Cross-session similarity was estimated by sampling 10 trials from each of two different sessions for the same image, averaging the response within each session, and computing the cosine similarity between the two session-specific response vectors. These estimates were averaged within each subject across resampling iterations and session pairs. Group-level differences between within-session and cross-session stability were then tested across subjects using non-parametric permutation tests.

For shuffle-based analyses, null distributions were constructed by disrupting the data structure required by the tested hypothesis, such as image identity, session pairing, or drift direction, while preserving the relevant marginal structure of the data. Observed statistics were then compared with the corresponding shuffled null distributions. The statistical tests used for comparing results from real drift condition and randomized direction drift (null) condition are paired t-test, with 95% confidence interval (CI).

For resampling-based significance tests, empirical null distributions were generated using 20,000 resamples unless otherwise stated. We computed p-values as the proportion of null statistics at least as extreme as the observed statistic, with the tail of the test chosen according to the stated hypothesis. When percentile intervals were reported, the lower and upper bounds were defined as the 2.5th and 97.5th percentiles of the corresponding resampled distribution.

## Supplemental Figures

**Figure S1:**
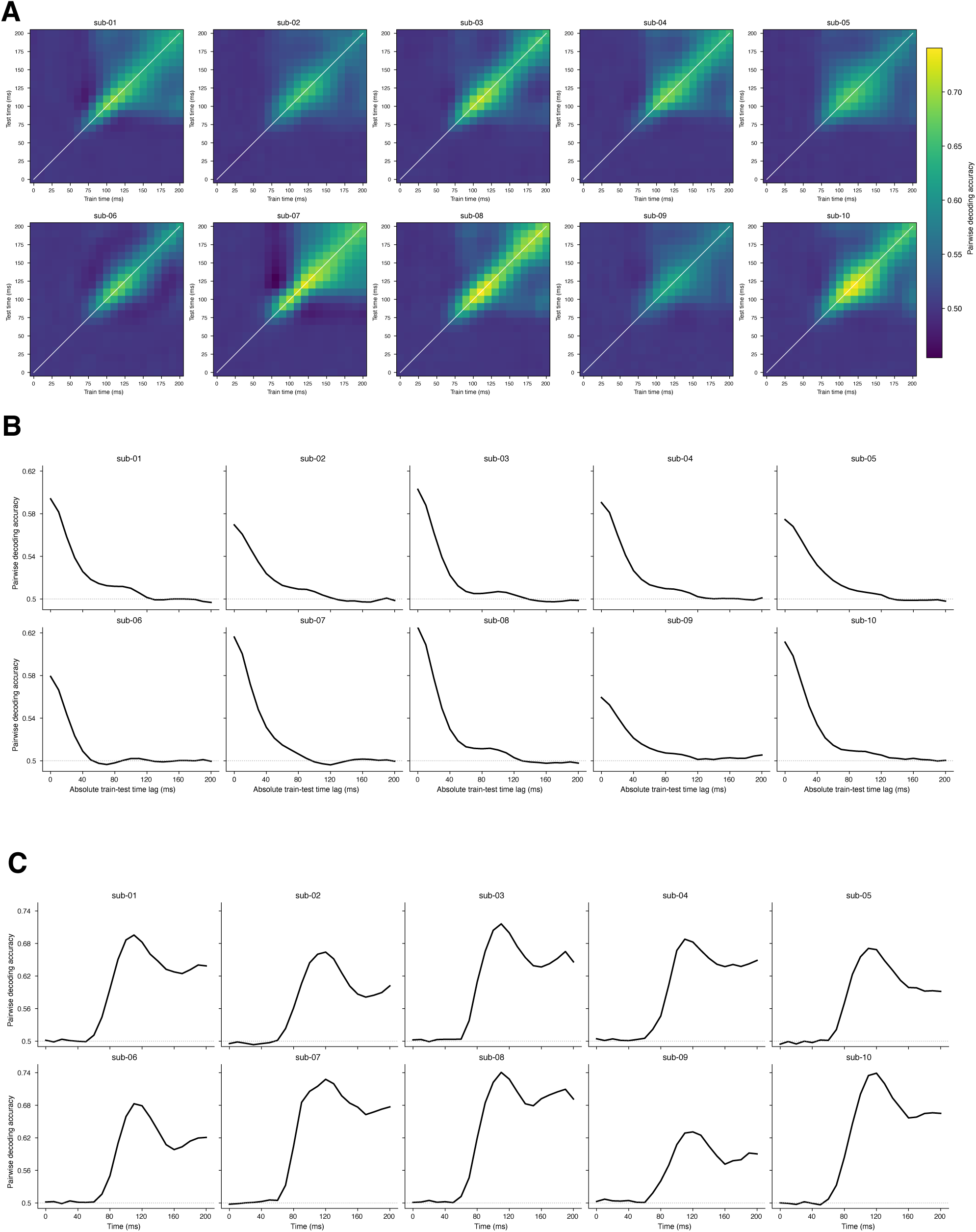
Decoding results for individual participants. (**A**) Cross-temporal pairwise image decoding. The x- and y-axes indicate training and testing time respectively, and each cell represents the decoding accuracy of a linear SVM trained at one time point and tested at another, averaged across image pairs. The diagonal therefore represents decoding when the training and testing time points were identical. Across participants, decoding accuracy remained near chance level until approximately 75 ms after stimulus onset and peaked at approximately 110 ms. Decoding was evaluated using five-fold cross-validation. For each pair of images, trials were randomly and evenly partitioned into five folds. When training and testing were performed at the same time point, four folds were used for training and the remaining fold was used for testing. For cross-temporal decoding, the classifier was trained in the same manner at one time point and evaluated using held-out trials at a different time point. This procedure was repeated until each fold had served as the test set. (**B**) Cross-temporal decoding accuracy as a function of temporal lag. The y-axis indicates decoding accuracy, and the x-axis indicates the (absolute value of the) temporal lag (Δ*t*) between the training and testing time points, ordered by the magnitude of the temporal interval. Across participants, decoding accuracy decreased as Δ*t* increased, indicating reduced generalization between increasingly separated time points. (**C**) Time course of within-session pairwise image decoding. The y-axis indicates five-fold cross-validated decoding accuracy at each time point (with decoder trained at the same time point), and the x-axis indicates time from 0 to 200 ms after stimulus onset. Decoding accuracy consistently peaked at approximately 110 ms across participants.

**Figure S2:**
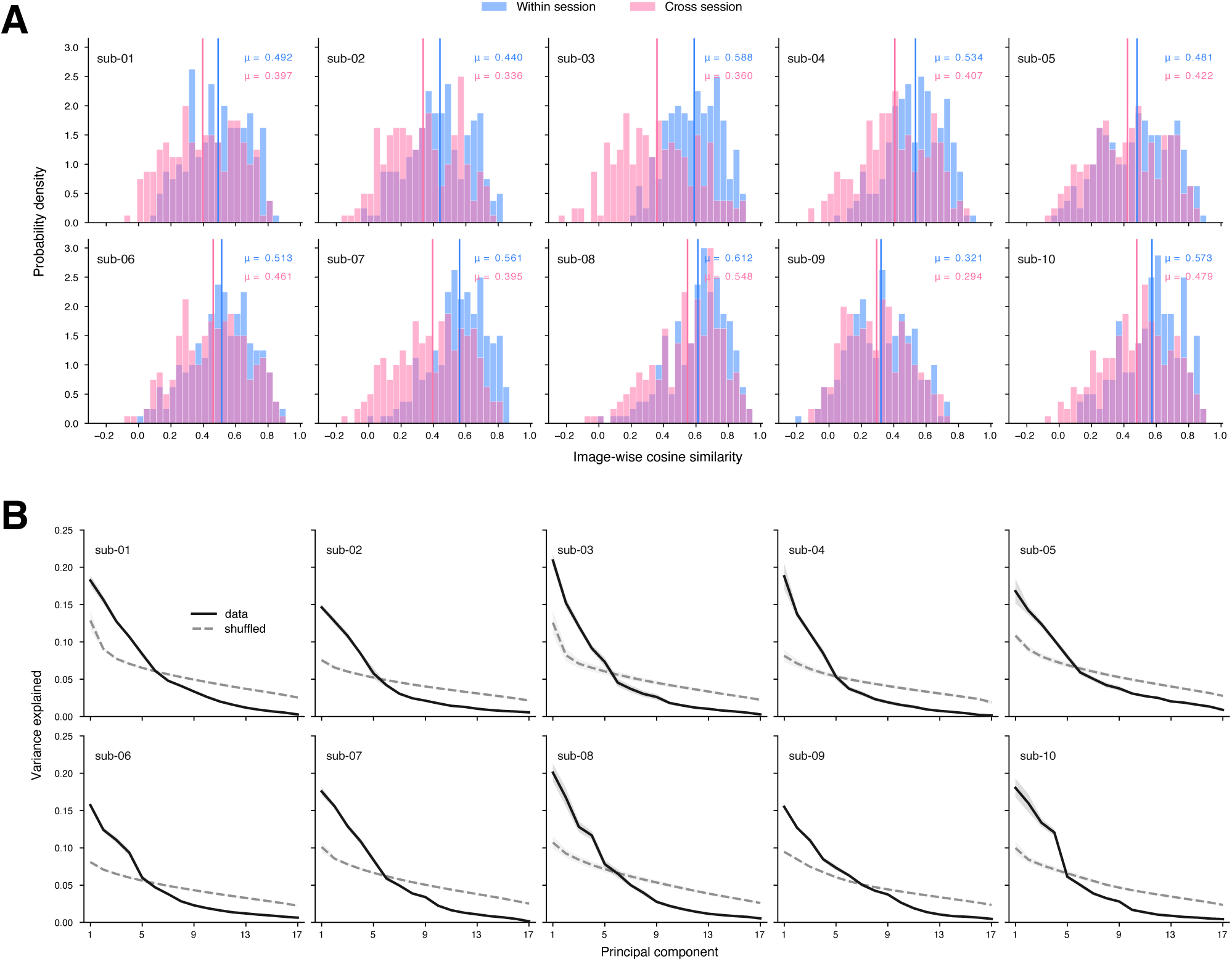
Within- and cross-session representational similarity and the structure of representational drift. (**A**) Distributions of same-image cosine similarities for within-session (blue) and cross-session (pink) comparisons, shown separately for each of 10 participants. Cosine similarity was computed between centered *K* = 17-dimensional EEG response vectors at 110 ms after stimulus onset. Comparisons used matched 10-trial averages, with disjoint trial halves for within-session comparisons. For each image, similarities were averaged over 2,000 sampling repetitions without replacement and all six session pairs, yielding 200 image-wise estimates per distribution. Vertical blue and pink lines indicate the means of the corresponding distributions.(**B**) Eigenvalue spectrum of the drift matrix compared with channel-shuffled control. The drift matrix was organized such that each row represented the cross-session drift vector for one image across EEG channels. To construct the shuffled control, image labels were independently permuted within each channel. Consequently, after shuffling, values within a given row no longer originated from the same image across channels. This procedure preserved the marginal distribution and variance of drift within each channel while disrupting the image-specific cross-channel structure of the drift vectors. The shuffled condition therefore provides a control for assessing whether the observed eigenspectrum reflects coordinated drift across channels.

**Figure S3:**
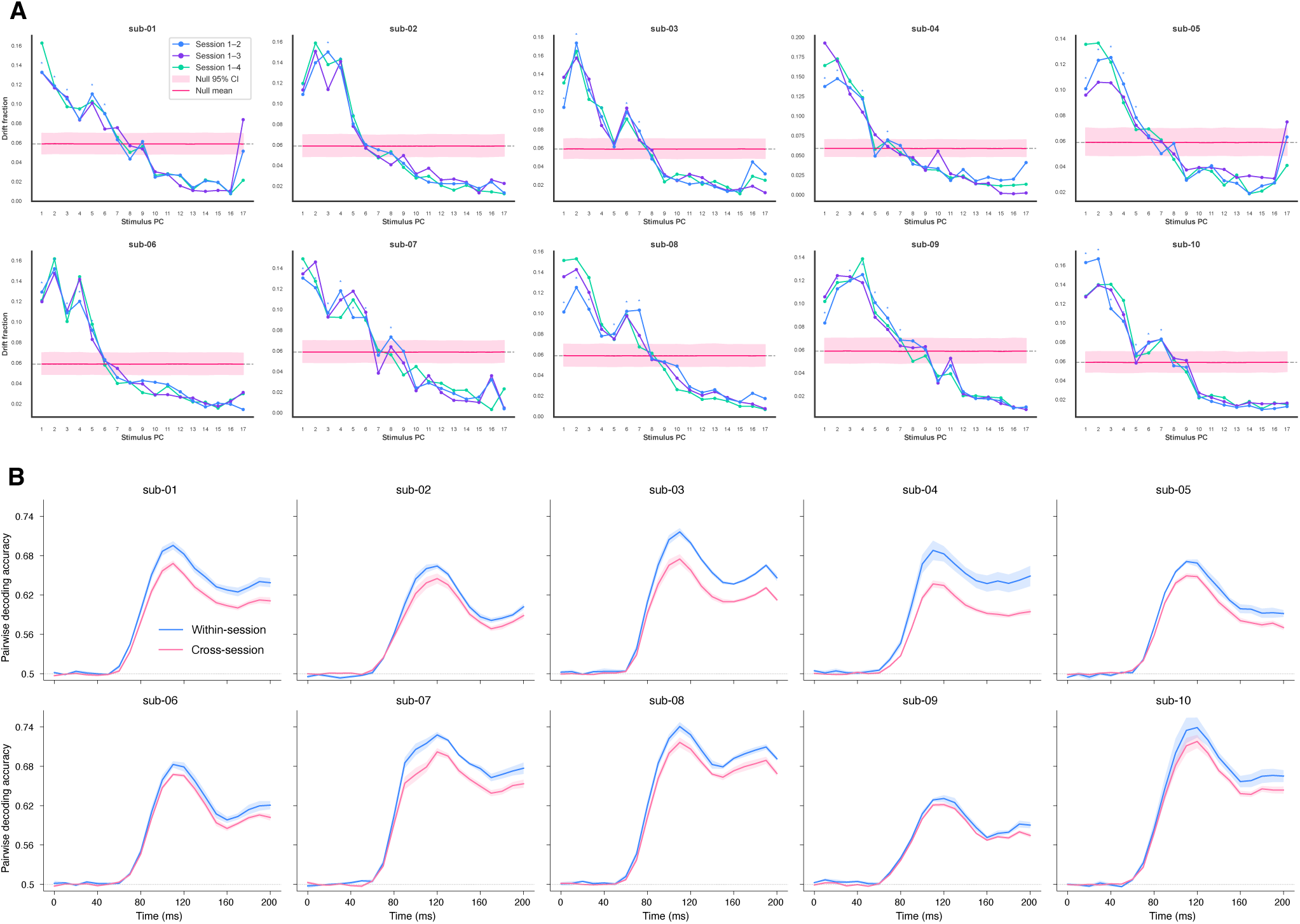
Representational drift is preferentially aligned with principal directions of the stimulus representation. (**A**) Drift is distributed nonuniformly across the principal components of the stimulus representation. The x-axis indicates individual principal components of the session 1 stimulus space, and the y-axis indicates the fraction of cross-session drift projected onto each component. Across session pairs, drift was preferentially concentrated along the leading principal components rather than being uniformly distributed across dimensions. Principal components explaining greater stimulus variance contained a larger fraction of drift than lower-variance components, indicating that drift preferentially occurred along dominant directions of the stimulus representation. Different colored lines represent different pairs of sessions. (**B**) Within-session pairwise image decoding accuracy was significantly higher than cross-session decoding accuracy across 10 participants beginning approximately 75 ms after stimulus onset. This within-versus cross-session decoding gap persisted after controlling for global session-level shifts in EEG activity, indicating that the reduction in cross-session decoding could not be explained solely by a global change in the measured population response.

**Figure S4:**
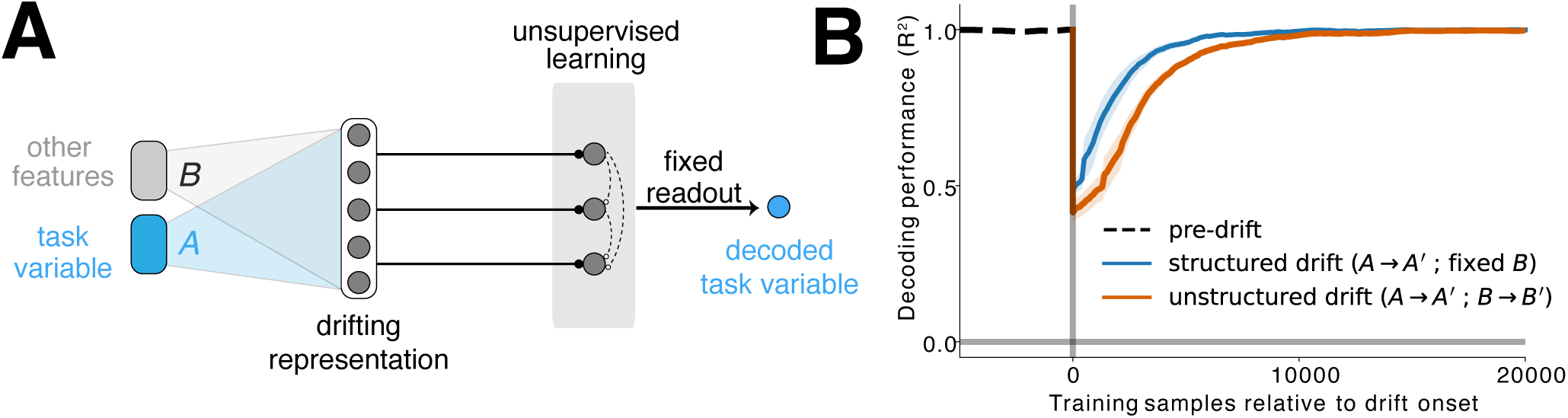
Faster recovery from structured than unstructured drift using Sanger’s learning rule. (**A**) We replaced the Hebbian/anti-Hebbian learning network with the multi-neuron Sanger model. As before, decoding of the task variable depended on the alignment between the learned network representation and the principal directions of the input representation. Learning consisted of two phases corresponding to the pre-drift and post-drift conditions. During phase 1, the network learned the principal components of the input representation, resulting in task-variable decoding performance approaching *R*^2^ = 1. The black dotted line indicates this pre-drift decoding baseline. After phase 1, we introduced either structured or unstructured representational drift. Structured drift was generated by an abrupt orthogonal rotation of the principal-component coordinates, whereas unstructured drift combined this rotation with an expansion of the effective dimensionality of the representation (Methods). To test whether ongoing unsupervised plasticity could compensate for these changes, the task-variable decoder learned during phase 1 was held fixed, while Sanger’s learning rule continued from the network state reached at the end of phase 1. (**B**) Both forms of drift initially reduced task-variable decoding performance. However, decoding recovered substantially faster after structured drift than after unstructured drift, requiring fewer additional training samples to return toward the pre-drift level.

## Notes

### Competing Interest Statement

The authors have declared no competing interest.

